# Infant Transmitted/Founder HIV-1 Viruses from Peripartum Transmission are Neutralization Resistant to Paired Maternal Plasma

**DOI:** 10.1101/183368

**Authors:** Amit Kumar, Claire E. P. Smith, Elena E. Giorgi, Joshua Eudailey, David R. Martinez, Karina Yusim, Ayooluwa O. Douglas, Lisa Stamper, Erin McGuire, Celia C. LaBranche, David C. Montefiori, Genevieve G. Fouda, Feng Gao, Sallie R. Permar

## Abstract

Despite extensive genetic diversity of HIV-1 in chronic infection, infant HIV-1 infection involves selective transmission of a single or few maternal virus variants. These transmitted/founder (T/F) variants are of particular interest, as a maternal or infant HIV vaccine should raise envelope (Env)-specific IgG responses capable of blocking this group of viruses. However, the maternal or infant factors that contribute to selection of infant T/F viruses are not well understood. In this study, we isolated HIV-1 *env* genes by single genome amplification from 16 mother-infant transmitting pairs from the U.S. pre-antiretroviral era Women Infant Transmission Study (WITS). Infant T/F and representative maternal non-transmitted Env variants from plasma were identified and used to generate pseudoviruses for paired maternal plasma neutralization sensitivity analysis. Eighteen out of 21 (85%) infant T/F Env pseudoviruses were neutralization resistant to paired maternal plasma. Yet, all infant T/F viruses were neutralization sensitive to a panel of HIV-1 broadly neutralizing antibodies and variably sensitive to heterologous plasma neutralizing antibodies. Moreover, infant T/F pseudoviruses were overall more neutralization resistant compared to maternal non-transmitted plasma variants (p=0.012). Altogether, our findings suggest that autologous neutralization of circulating viruses by maternal plasma antibodies select for neutralization-resistant viruses that initiate peripartum transmission, raising the spector that enhancement of this response at the end of pregnancy could further reduce infant HIV infection risk.

**Author Summary:** Mother to child transmission (MTCT) of HIV-1 can occur during pregnancy (*in utero*), at the time of delivery (peripartum) or by breastfeeding (postpartum). With the availability of anti-retroviral therapy (ART), rate of MTCT of HIV-1 have been significantly lowered. However, significant implementation challenges remains in resource-poor areas, making it difficult to eliminate pediatric HIV. An improved understanding of the viral population (escape variants from autologous neutralizing antibodies) that lead to infection of infants at time of transmission will help in designing immune interventions to reduce vertical HIV-1 transmission. Here, we selected 16 HIV-1-infected mother-infant pairs from WITS cohort (from pre anti-retroviral era), where infants became infected peripartum. HIV-1 *env* gene sequences were obtained by the single genome amplification method. The sensitivity of these infant Env pseudoviruses against paired maternal plasma and a panel of broadly neutralizing monoclonal antibodies (bNAbs) was analyzed. We demonstrated that the infant T/F viruses were more resistant against maternal plasma than non-transmitted maternal variants, but sensitive to most (bNAbs). Signature sequence analysis of infant T/F and non-transmitted maternal variants revealed the potential importance of V3 and MPER region for resistance against to paired maternal plasma. These findings provide insights for the design of maternal immunization strategies to enhance neutralizing antibodies that target V3 region of autologous virus populations, which could work synergistically with maternal ARVs to further reduce the rate of peripartum HIV-1 transmission.

## Introduction

Despite the wide success of antiretroviral therapy (ART) in lowering mother-to-child transmission (MTCT) risk of HIV-1 below 2%, each year more than 150,000 children become infected worldwide [1]. Even if 90% maternal ART coverage is reached, approximately 138,000 infant HIV-1 infections will still occur annually [2, 3] due to factors that include: drug non-adherence, breakthrough infections, development of drug resistant viral strains, late presentation of pregnant women to clinical care, and acute infection during late pregnancy or breastfeeding. Vertical transmission of HIV can occur through three distinct modes: antepartum (*in utero)*, peripartum (around the time of delivery), or postpartum (via breastfeeding). Interestingly, only 30-40% of infants born to HIV infected mothers acquire HIV-1 in the absence of ART[4]. Thus, maternal factors, such as maternal Env-specific antibodies, may contribute to protecting infants from HIV infection. Maternal factors that are associated with HIV transmission risk include: low maternal peripheral CD4+ T cell count, and high maternal plasma viral load, delivery mode, and infant gestational age [5-7]. Yet, the role of maternal Env-specific antibody responses and their association with reduced MTCT risk still remains unclear. Previous studies have reported an association between the magnitude of maternal antibody responses and reduced risk of MTCT [8-10]. However, this association has not been universally observed [11-15]. Moreover, it has been observed that variants transmitted to infants can be resistant to neutralization by maternal plasma [16], although other studies have failed to replicate these observations [17-19]. These conflicting results may be due to the small number of subjects included in these studies and study designs that inconsistently control for viral and host factors known to impact transmission risk, such as maternal peripheral CD4+ T cell counts, plasma viral load, non-identification of T/F viruses, and ART use. Moreover, isolation of autologous viruses from a large cohort of HIV-infected, transmitting mothers for assessment of the impact of maternal plasma neutralization activity against her own viruses has not to our knowledge, been investigated. Thus, despite considerable effort, it remains unclear whether maternal antibody responses impact the risk of vertical transmission of HIV.

We recently completed a maternal humoral immune correlates of protection analysis to identify maternal humoral immune responses associated with protection against peripartum HIV infection using samples from the US-based Women and Infants Transmission (WITS) study [20]. The WITS cohort was enrolled prior to the availability of ART prophylaxis as the clinical standard of care in HIV-infected pregnant mothers and their infants, thereby eliminating the strong impact of ART on vertical HIV transmission risk and outcome [21, 22]. Additionally, we controlled for established maternal and infant risk factors associated with vertical transmission, including maternal peripheral CD4+ T cell count, maternal plasma HIV-1 viral load, infant gestational age, and delivery mode by propensity score matching of transmitting and non-transmitting women. The results of this immune correlate analysis indicated an association between high levels of maternal antibodies against the HIV-1 Env glycoprotein third variable loop (V3) and reduced MTCT risk [20]. In addition, and more surprisingly, the ability of maternal plasma to neutralize tier 1 viruses (easy-to-neutralize), but not tier 2 (difficult to neutralize) viruses, also predicted decreased risk of vertical transmission of HIV-1. Yet, vertically transmitted HIV variants have been characterized as more difficult to neutralize tier 2-like variants [17, 23-26]. Thus, it was surprising that tier 1 virus neutralizing antibodies were associated with decreased transmission risk. More interestingly, maternal V3-specific monoclonal IgG antibodies isolated from a non-transmitting mother neutralized a large proportion of maternal autologous viruses isolated from her plasma[20], leading to the conclusion that maternal V3-specific non-broadly neutralizing antibodies, which were previously thought to be ineffective at preventing HIV-1 transmission, might indeed play a role in preventing MTCT. In fact, Moody et.al [27] showed that V3 and CD4 binding site (CD4bs) specific monoclonal antibodies isolated from non-pregnant chronically HIV-infected individuals could also neutralize a large proportion of autologous circulating viruses isolated from plasma. These V3 and CD4bs-specific autologous virus-neutralizing mAbs exhibited tier 1 neutralization activity but limited heterologous tier 2 virus neutralization, suggesting that measurement of tier 2 heterologous virus neutralization potency of mAbs or plasma does not predict autologous virus neutralization capacity.

In contrast to the extensive genetic diversity of HIV-1 variants in a chronically infected host, acute HIV infection in both heterosexual and vertical routes are characterized by a homogeneous viral population [17, 18, 28-31]. This viral genetic bottleneck suggests the selective transmission of a single or homogeneous group of viruses [4]. In the setting of MTCT, maternal or infant immunologic and virologic factors that that drive the selective transmission of one or a few HIV variants are not established [32]. As maternal viruses co-circulate with maternal HIV Env-specific antibodies, it is possible that maternal antibodies play a role in selecting maternal escape viruses that may initiate infection in the infant. Therefore, studying unique features of infant T/F viruses and their neutralization-sensitivity determinants to maternal autologous virus neutralizing antibodies may provide insights of the molecular events that lead to virus escape from maternal humoral responses.

The use of broadly neutralizing antibodies as a treatment and/or prevention strategy is currently being explored in adult and infant clinical trials [33, 34]. Among the new generation of bNAbs, VRC01 (antibody recognizing CD4bs region) has been able to neutralize about 80% of diverse HIV-1 strains [35, 36]. This has lead to studies of VRC01 impact on HIV-1 infection in adults and infants when infused passively, with a phase I study of pharmacokinetics and safety of VRC01 in HIV-exposed newborns currently underway [37]. However, the susceptibility of infant T/F viruses to bNAbs like VRC01 does not seem to define infant T/F viruses from maternal non-transmitted viruses [17] and administration of a bNAb to chronically-infected mothers is likely to lead to rapid development of resistant viruses [34, 38]. Thus, defining the role of autologous neutralization in MTCT is critical to establishing the utility of active maternal vaccination to further reduce and eliminate infant HIV infections.

In this study, we characterized maternal non-transmitting and infant T/F viruses from 16 HIV-1 clade B infected peripartum transmission mother-infant pairs from the WITS cohort and defined the role of concurrent maternal autologous virus neutralizing antibodies in selecting for infant T/F viruses. We sought to define if neutralization resistance to paired maternal plasma was a defining feature of infant T/F viruses compared to other circulating non-transmitted maternal variants, which will inform the development of maternal or infant vaccination strategies to further reduced MTCT risk to achieve an HIV-free generation.

## Results

### Maternal and infant sample characteristics

We selected sixteen, HIV-1 infected mother-infant transmission pairs from the WITS cohort that met the inclusion criteria of peripartum transmission (infants tested negative for HIV-1 infection at birth by HIV-1 DNA PCR, yet had HIV-1 DNA detectable at one week of age or older; see Table1). These HIV-exposed infants had not reportedly been breastfed [22]. Infant plasma samples available for sequencing were between 16 - 74 days of age. Five HIV-infected infants with heterogeneous virus populations were excluded from the study due to our inability to confidently infer the infant T/F virus. The maternal plasma viral load of the selected transmitting women ranged from 4,104 to 3,68,471 copies/mL, and peripheral blood CD4^+^ T-cell counts ranged from 107 to 760 cell/mm^3^. Infant plasma viral loads varied between 11,110 and 2,042,124 copies/mL, and CD4^+^ T-cell counts were between 1,872 and 7,628 cell/mm^3^. All infants were born via vaginal delivery except for three infants (100014, 100155, 100307) born via cesarean section, thus potentially representing late *in utero* transmission. Four infants (100014, 100155, 100307, 102149) were born prematurely (31, 34, and 36 weeks respectively), and the remaining infants were born between 37 and 40 weeks of gestation.

### Characterization of the complete *env* gene sequences from paired mother-infant plasma

A total of 463 *env* genes were obtained from 16 maternal plasma samples (collected at time of delivery) from transmitting mothers as previously described [24]. Paired infant plasma samples were used to obtain 465 *env* gene sequences (Table 2). Neighbor-joining phylogenetic trees and highlighter plots of the *env* sequences from each infant were used to define infant T/F viruses. These analyses showed within-lineage low diversity populations in infant Env isolates and chronic-like diversity in maternal *env* sequences (Fig.1, A-C and Fig. S1 A-B). In 6 out of 16 (37%) infants we detected 2 or 3 (in case of 100002) genetically distinct T/F variants, one of which was present at higher frequency (primary T/F), while the second one was present at lower frequency (secondary T/F). In the other 10 infants demonstrated we observed only one T/F virus (67%). With the exception of two infants, all of our samples had over 20 infant sequences, giving us a 90% confidence that we were able to sample all variants with a population frequency of at least 10%. For the two infant samples for which we only had 15 and 18 sequences respectively, we were 90% confident that we were able to sample all variants with a population frequency of 15% or more [24]. Using an algorithm described in the Methods, out of these 463 maternal *env* variants, we selected 134 SGA variants for *env* pseudovirus production (5-12 per mother) to represent the *env* genetic diversity found in the plasma of each transmitting mother at the time of delivery.

**Fig. 1.**
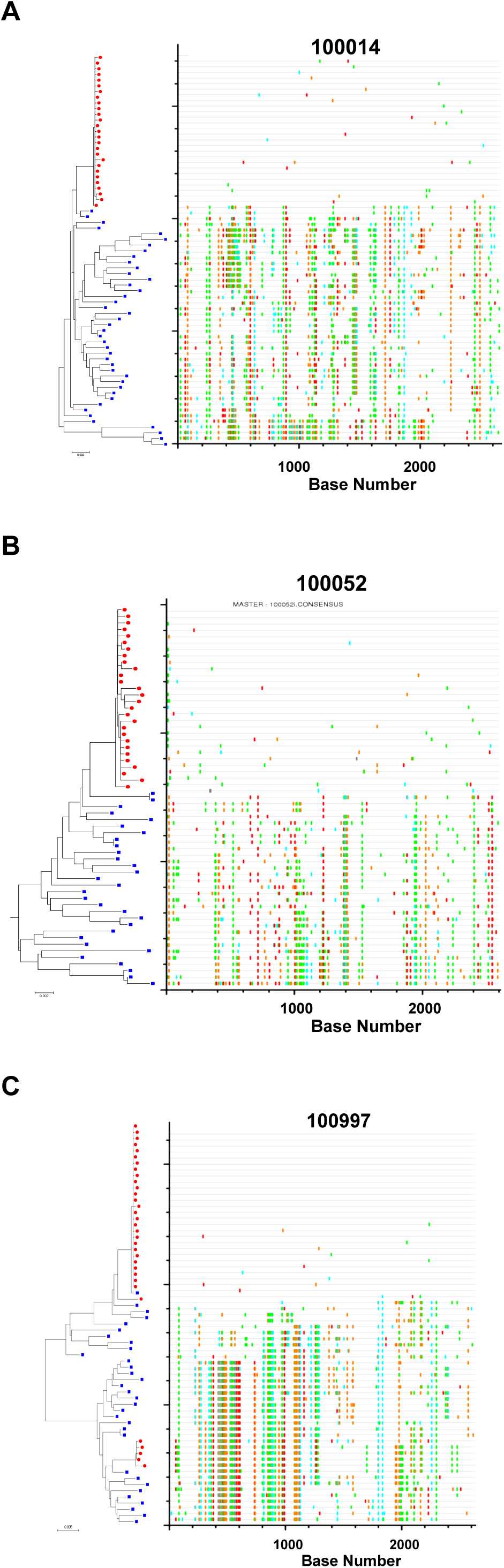
Highlighter plots of maternal and infant viruses. Example of mother-infant pair where the infant was infected with a single T/F virus and no evidence of evolutionary selection in the infant (3/16 infants) (A). Example of mother infant with evidence of evolutionary selection in the infant (B), and Example of mother infant pair where the infant was infected by two distinct T/F viruses (6/16) infants (C). Individual infant and maternal viruses are represented by red dots and blue squares, respectively, on the tree. Colored hash marks on each highlighter plot represent nucleotide differences as compared with the infant consensus sequence at the top and are color-coded according to nucleotide.

### Confirming the timing of infant HIV-1 infection

As the infants were selected for peripartum transmission, their age at sampling (in days) was also the post-infection time. To confirm the time of infection, all infant alignments were analyzed using the LANL Poisson Fitter tool [39]. For infants that had more than one T/F, only the sequences in the major T/F lineage were used for this analysis. When recombinants and APOBEC enrichment were detected, the timing was calculated after removing recombinants and/or positions enriched for hypermutation [24, 39].

All but one infant (102605) yielded a good Poisson fit, indicating that the amount of diversity found in these samples was compatible with a random accumulation of mutations as observed in acute infections. Four infants had detected recombinants and 6 infants yielded a good Poisson fit after removing positions enriched for hypermutation (Table 2). The time since the most common ancestor was consistent with transmission at delivery in 9 out of 16 pairs (57%), within the 95% confidence interval of the Poisson Fitter time estimate (Table 2). For three infants, Poisson Fitter estimated the time since infection to be younger than the reported infant age, and for one the infection showed more diversity than expected by the reported infant age. Discrepancies between actual vs. predicted transmission timing could be due to a number of factors, including late *in utero* infection, postpartum infection from unreported breastfeeding, and the model being designed to evaluate an adult rather than infant HIV-1 evolution, which could gather mutations more rapidly due to more robust T cell responses.

Infant 102605 *env* SGAs did not yield a good Poisson fit due to non-random accumulation of non-synonymous mutations (which breaks the model assumption of random accumulation of mutations) at HXB2 positions 752-754 (Fig. S2). We looked at this region in the LANL immunology database (https://www.hiv.lanl.gov/content/immunology/index), and found five different human CTL epitopes that have been documented in the literature, confirming that the non-random mutations found in infant 102605 were likely due to selection pressure by T cell responses.

### Neutralization sensitivity and tier classification of the infant T/F viruses

Twenty-one infant T/F *env* amplicons including 16 primary T/Fs and 5 secondary T/Fs were used to generate pseudoviruses and their neutralization sensitivity to paired maternal plasma and a panel of bNAbs was assessed. None of the mothers with the exception of two (100014 and 100504) had non-specific neutralization activity as assessed by neutralization activity against a murine leukemia virus (MLV). Eighteen out of 21 infant T/F Env pseudoviruses (86%) were resistant to paired maternal serum (ID_50_<40). Sensitivity of 2 Infant T/F pseudoviruses’ (100014 and 100504) against paired maternal plasma could not be determined with confidence due to higher plasma reactivity against MLV. T/F virus of infant 100046 was sensitive against paired maternal plasma (Fig. 2).

**Fig. 2:**
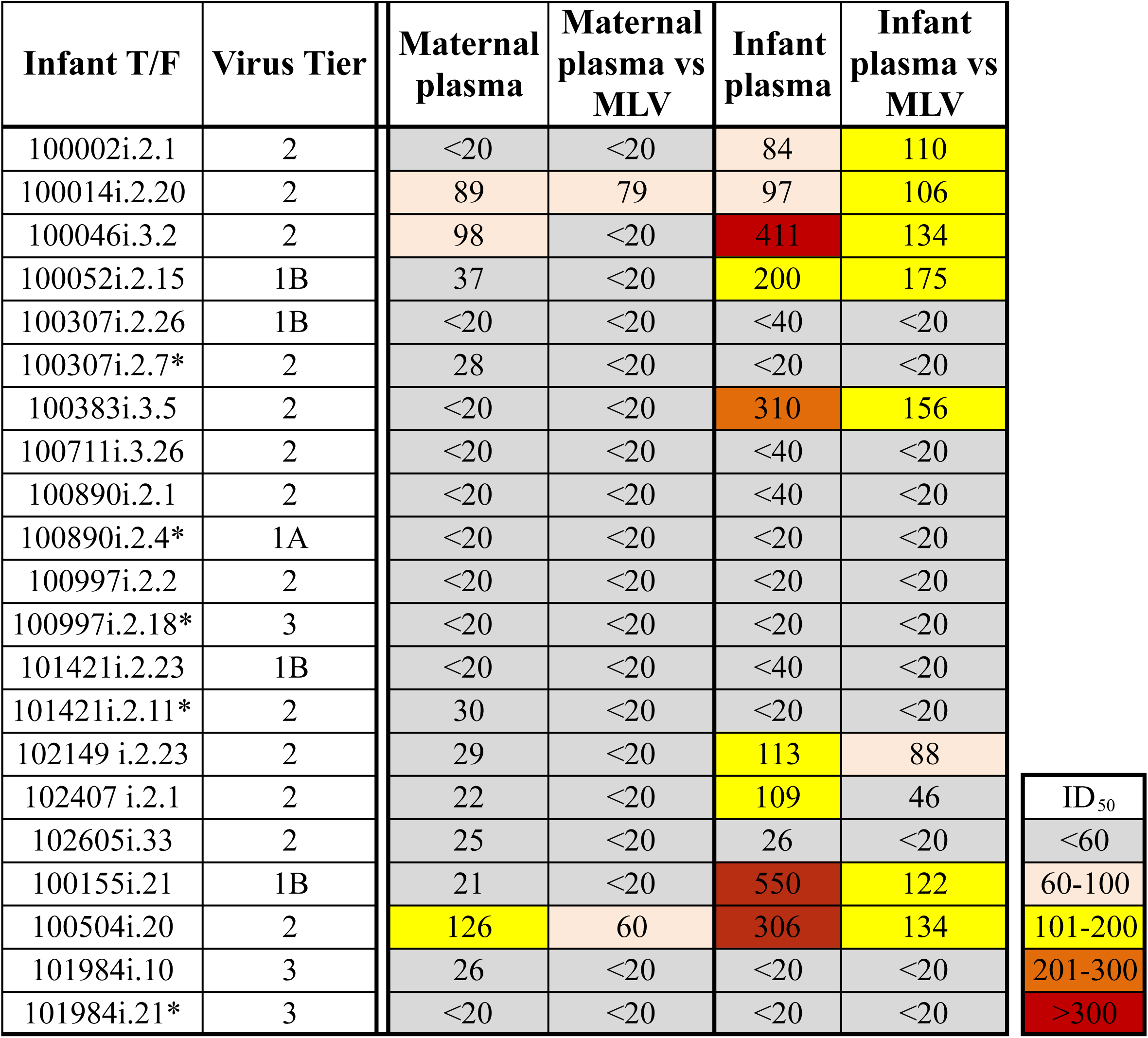
Tier phenotyping and neutralization sensitivity of Infant T/F viruses to paired maternal plasma and heterologous broadly neutralizing antibodies (bNAbs). Dark colors represent easily neutralized viruses. Second T/F viruses are marked with a star (*).

When Infant T/F pseudoviruses were tested against the autologous plasma, only 2 infants T/F (100046 and 100155) showed some sensitivity as per our criteria (ID_50_> 3X that of MLV) while others were completely resistant. Some infant T/F pseudoviruses did show sensitivity against their own plasma but were not considered as sensitive due to high reactivity against MLV.

To determine whether these infant T/F *env* variants were globally resistant to heterologous plasma neutralization, we performed neutralization tier phenotyping using a standardized panel of heterologous plasma of HIV-1 infected individuals [40]. Thirteen (62%) of 21 infant T/F Env pseudoviruses tested were classified as tier 2 neutralization phenotype while 3 (14%) of 21 were classified as tier 3 neutralization phenotype, as expected for infant T/F viruses (Fig. 2). Remarkably, the remaining 5 (24%) tested were classified as the easier-to-neutralize tier 1b or tier 1a sensitivity, possibly because that these variants were uniquely resistant to their paired maternal plasma (Fig. 2).

In contrast to the relative resistance to paired maternal plasma neutralization, all the infant T/F viruses were relatively sensitive to second-generation HIV-1 broadly neutralizing antibodies, such as VRC-01, (IC_50_ range 0.12-5.0 μg/ml), PGT121 (IC_50_ range 0.01-0.13 μg/ml), NIH 45-46 (IC_50_ range 0.01-0.28 μg/ml) and first generation bNAb 10E8 (IC_50_ range 0.05-0.76 μg/ml) (Fig. 3 and Fig. S3). Not surprisingly, the infant T/F Env pseudoviruses were less neutralization sensitive to the less potent first generation broadly neutralizing antibodies b12 (IC_50_ range 2.97-25 μg/ml), 4E10 (IC_50_ range 1.31-18.73 μg/ml) and 2F5 (IC_50_ range 1.01-19.09 μg/ml) (Fig. 3). Importantly, all the infant T/F viruses were neutralization sensitive to VRC-01, a bNAb currently being evaluated in clinical trials for use in HIV exposed infants. However, V3 glycan-specific bNAbs, which clustered together in neutralization sensitivity, and NIH45-46 mediated the most neutralization breadth and potency against the infant T/F viruses (Fig. 3).

**Fig. 3.**
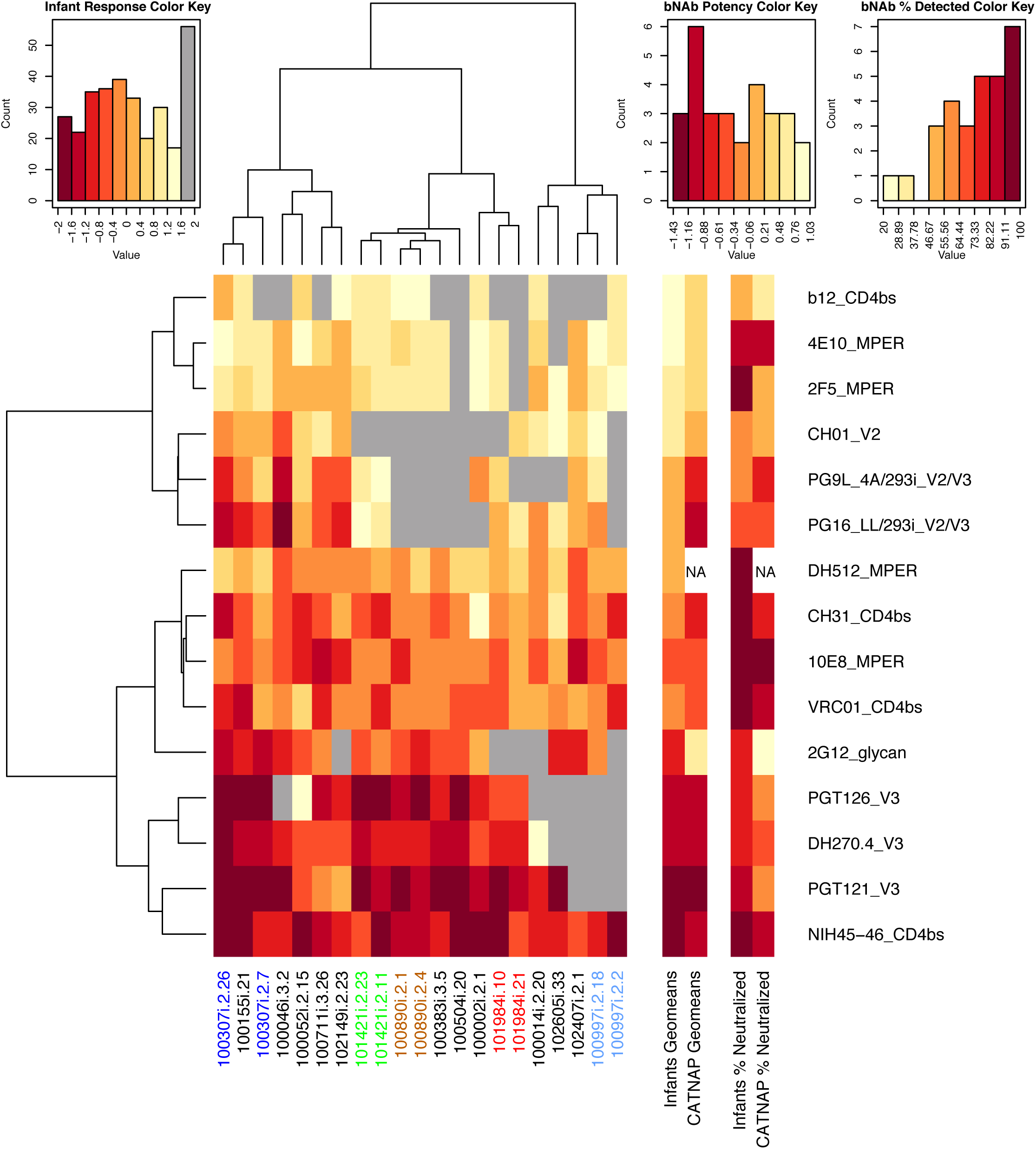
Infant neutralization sensitivity against a panel of bNAbs. The strength of the responses is color coded from dark to light where darker reds indicate stronger responses. Aquamarine indicates absence of response. T/F viruses from the same infant are labeled using the same color (left columns). The potency (geometric mean of responses) and breadth (% neutralized) of the infant viruses to each bNAb were compared with the geometric means of the bNAb’s potency and breadth obtained from published studies using the LANL repository CATNAP (columns on the right). The most potent neutralization against infant viruses was mediated by V3 glycan bNAbs and CD4 binding site specific NIH45-46.

We next calculated the geometric means of both the breadth and potency of the panel of bNAbs against the infant T/F viruses and compared with their potency against other HIV variants as documented in CATNAP (Compile, Analyze and Tally NAb Panels) [41], the Los Alamos National Laboratory (LANL) interface that collects all published immunological data. In general, potency and breadth of the bNAbs against infant T/F viruses followed the potency and breadth calculated in CATNAP (p=0.013 and 0.02 respectively, Spearman correlation test), with one exception: bNAb 2G12 displayed more potent and broad responses in the infants than in the CATNAP collective data (Fig. 3).

### Neutralization sensitivity of non-transmitted maternal *env* variants compared to infant T/F *env* variants

Pseudoviruses were prepared from a total of 134 non-transmitted maternal *env* variants using the promoter PCR method [42] and assessed for neutralization sensitivity against paired maternal plasma, including the isolated maternal non-transmitted variant that was most closely related to the infant T/F variant. Variable neutralization sensitivity to paired maternal plasma was observed in non-transmitted maternal variants, with some of the variants exhibited neutralization sensitivity while others showed complete neutralization resistance (Fig. 4). Comparison of neutralization sensitivity between infant T/F Env variants and the identified closest maternal variant within each mother-infant pair revealed no consistent pattern. Over a 2-fold increase in sensitivity was observed for Env pseudoviruses of non-transmitted maternal variants that were most closely related to infant T/F for 6 infants (100002, 100307, 100052, 102149, 102407 and 102605). In contrast, infant T/F Env pseudoviruses from 3 infants (100014, 100046 and 100504) were more sensitive to maternal plasma than their most closely related maternal variants.

**Fig. 4.**
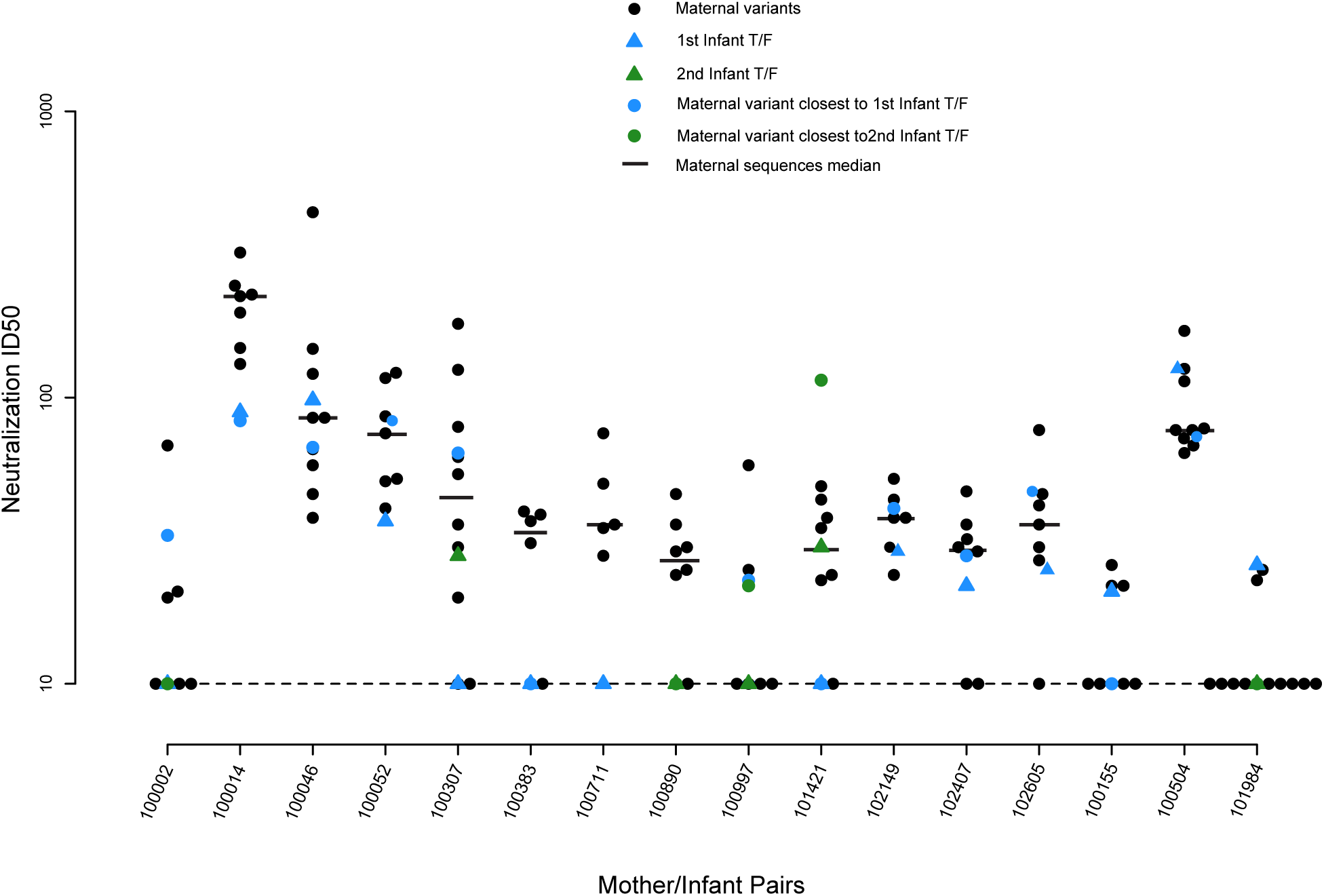
Neutralization sensitivity of maternal and infant viruses to paired maternal plasma at delivery. Sensitivity of maternal non-transmitted variants (black dots), infant T/F variants (blue and green triangles) and the closest maternal variant (blue and green dots) to the infant T/F variant against autologous maternal sera. Sequences were selected following an algorithm to represent different motifs that diverge from the infant T/F (see Methods). Black horizontal lines represent the median of the ID_50_ of maternal and infant sequences. The dashed line represents the detection threshold.

Yet, the infant T/F viruses were generally more resistant to the maternal plasma at delivery than the non-transmitted viruses from mothers within each maternal-infant pair, with the exception of 100046. However, since there were only 1 or 2 T/F viruses in each infant, we could not perform statistical analysis to determine if the differences are statistically significant within each pair. To determine whether infant T/F viruses were overall more resistant to maternal plasma than the paired non-transmitted maternal viruses, we employed a 1-sided permutation test to compare the neutralization sensitivity of maternal non-transmitted variants to the infant T/F Env variants. Remarkably, infant T/F Env variants were overall significantly more resistant to paired maternal plasma collected at delivery than non-transmitted maternal Env variants (p=0.01). Even when excluding the mother-infant pairs with high MLV neutralization (100014 and 100504), the infant T/F Env variants remained more resistant to neutralization than non-transmitted maternal variants (p=0.005).

To assess whether any particular epitope-specific neutralization sensitivity was distinct in infant T/Fs compared to matched maternal variants, we determined the neutralization sensitivity of 4 bNAbs targeting distinct vulnerable epitopes on HIV-1 Env: VRC-01 (CD4bs-specific) (VRC-01), PG9 (V2 glycan-specific), DH429 (V3 glycan-specific), and DH512 (membrane proximal external region – MPER-specific) (Fig. 5). We used the same 1-sided permutation test described above to assess for differences in neutralization sensitivity to these bNAbs in infant T/Fs vs non-transmitted maternal sequences. Interestingly, we found that infant T/F viruses were significantly more resistant to DH512 (MPER-specific) compared to non-transmitted maternal sequences (p = 0.025 by 1-sided permutation test; p=0.045 when excluding the two mothers with non-specific neutralization), while all other comparisons yielded no statistical significance (Fig. 5).

**Fig. 5.**
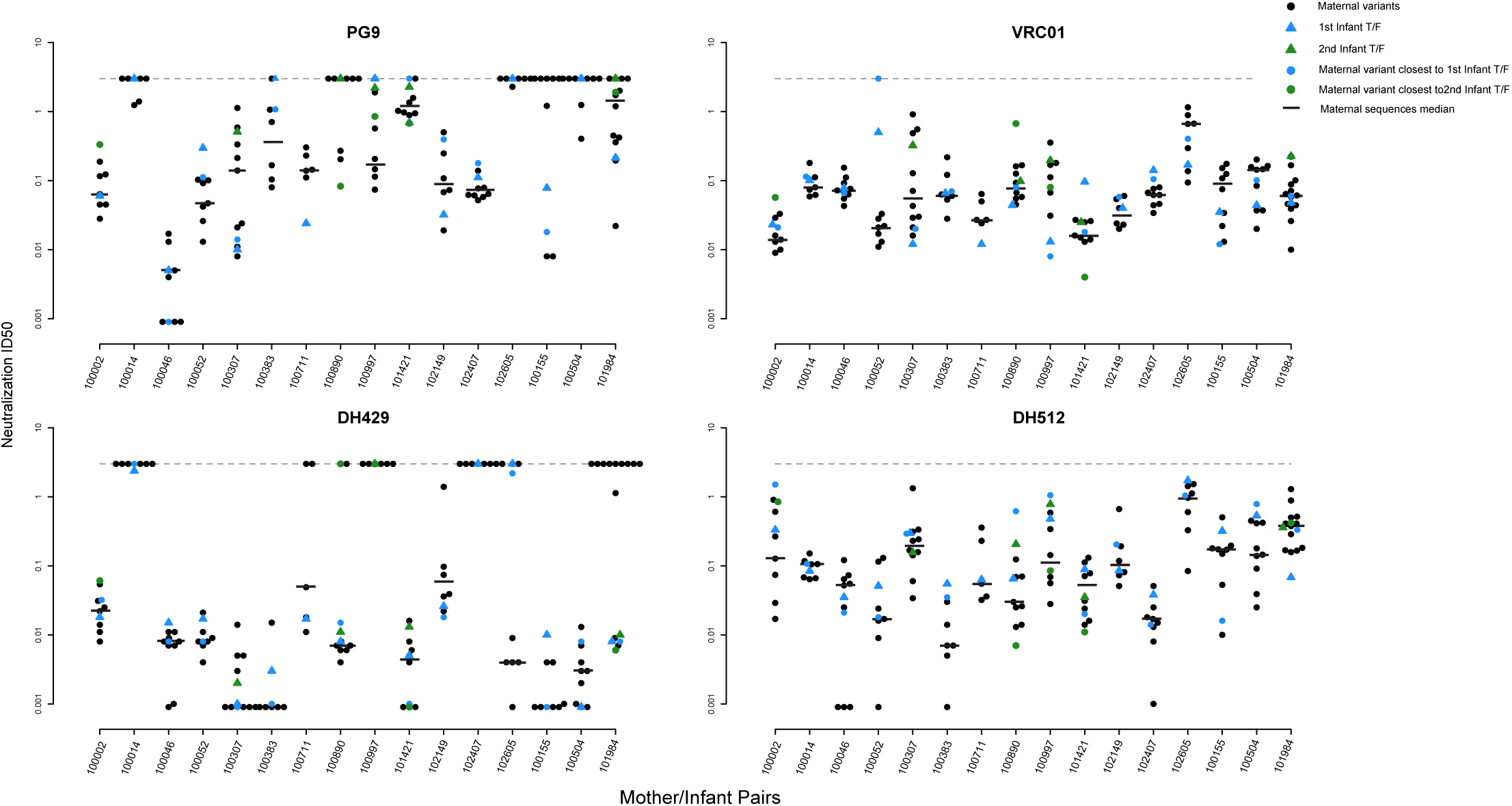
Neutralization sensitivity of maternal and infant viruses to paired maternal plasma and an MPER specific bNAb with and without identified signature sequences within MPER. IC_50_ of maternal non-transmitted variants (black dots), infant T/F variants (blue and green triangles) and the closest maternal variant (blue and green dots) to the 4 antibodies PG9, VRC01, DH429, and DH512. Black horizontal lines represent the median of the IC_50_ of maternal and infant sequences. Dashed lines represent the detection threshold.

### Signature sequence analysis of infant T/F variants to predict neutralization resistance

Because DH512 binds to the MPER region, we investigated the amino acid positions within this epitope (positions 662-683) and identified 4 positions that were either associated with higher maternal plasma neutralization (position 662, amino acid A, K, Q, or S were significantly more resistant than the wild type E, p=9.9e-05, 1-sided permutation test), lower DH512 IC_50_ (position 667 and 676, p=9.9e-04 and 0.003, respectively by 1-sided permutation test), or both (position 683, amino acid R was significantly more resistant than the wild type K (most frequent AA at this position), p<1e-04 by 1-sided permutation test; Fig. 7). However, when we looked at the Env sequences in individual mother-infant pairs, these amino acid residues that associated with DH512 neutralization resistance were equally distributed across non-transmitted maternal sequences and infant T/F viruses. Therefore, we could not to determine if T/F variants enriched for neutralization resistance-conferring amino acids were more apt to be transmitted compared to non-transmitted maternal variants. This could be due to the low sequence number within pairs, or it could suggest that the wild type amino acids at these positions are associated with DH512 neutralization resistance but not necessarily transmission.

**Fig. 6.**
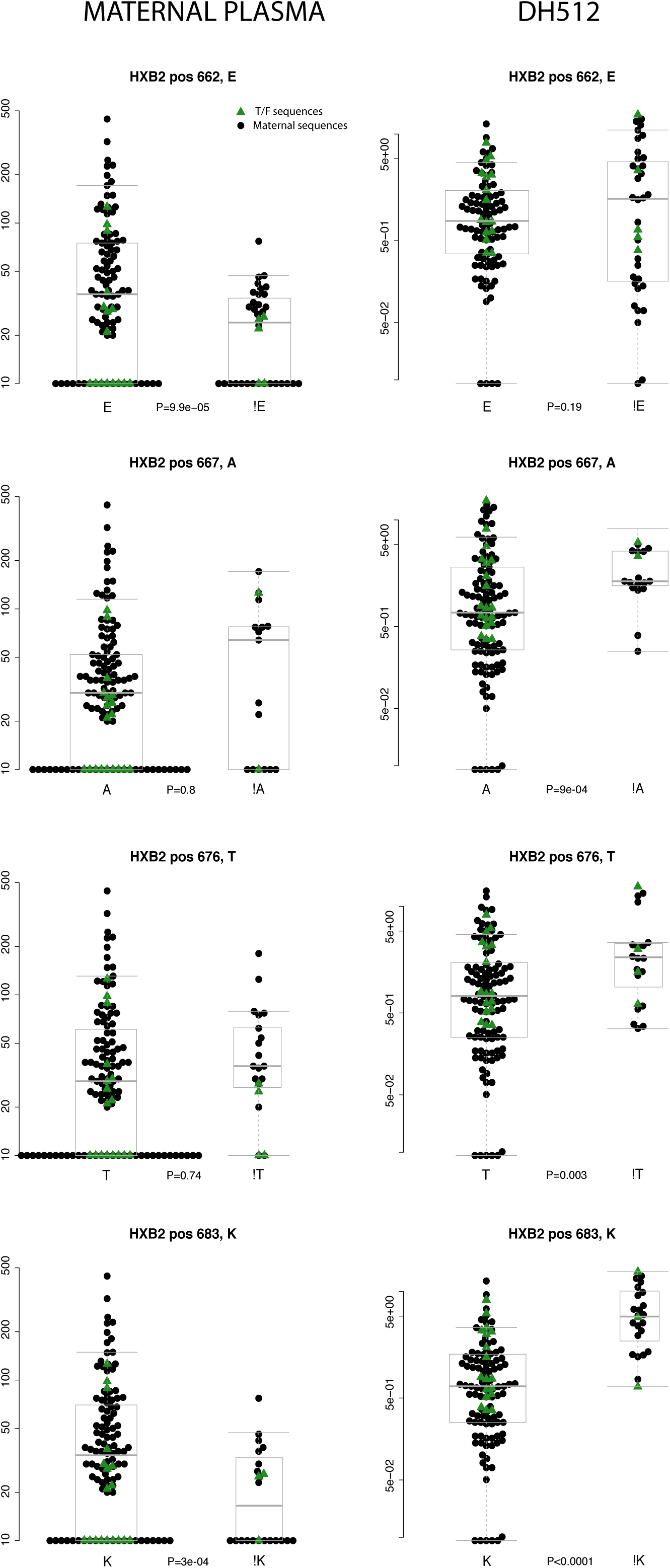
Comparison of neutralization sensitivity of maternal and infant viruses to epitope specific bNAbs. Comparison of paired maternal plasma and MPER specific bNAb DH512 maternal non-transmitted sequences (black dots) and infant T/F sequences (green triangles) between sequences that carry the wildtype amino acid at HXB2 positions 662, 676, 676, and 683, compared with those that carry a mutant. These four positions were chosen because they yielded a significant association with either maternal plasma responses (i.e. sequences carrying the mutant were statistically significantly more resistant to maternal plasma) or with DH512 responses (i.e. sequences carrying the mutant were statistically significantly more resistant to DH512) in the MPER epitope. P-values were obtained using a 1-sided permutation test (see Methods). Gray boxes represent median and quartiles of the responses.

**Fig. 7.**
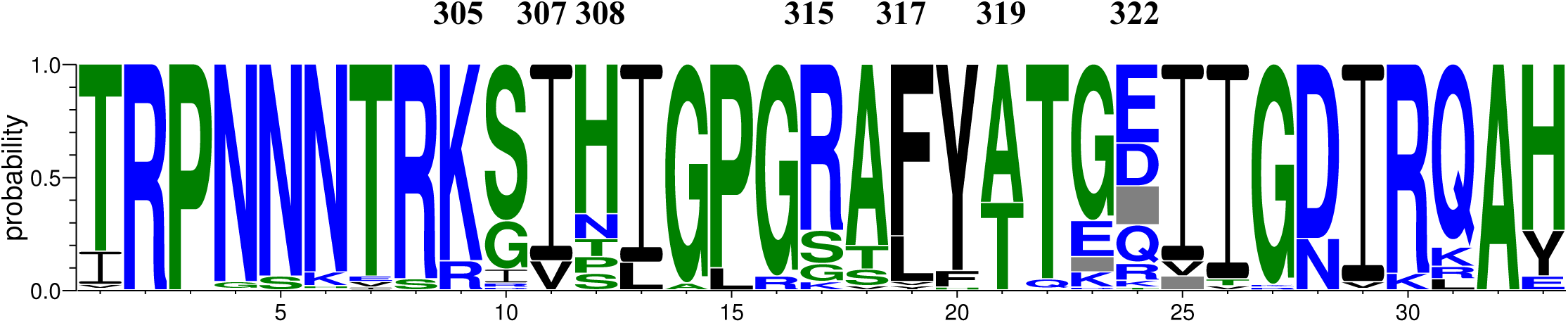
Frequency plot of the V3 region sequences of mother and Infant T/F viruses. Weblogo plot showing frequency of amino acids in the V3 region was made using the AnalyzAlign tool from the LANL website. Positions were numbered based on HXB2 amino acid sequence. Amino acid positions identified in the signature sequence analysis are marked at the top of the plot.

### V3 loop amino acid signature sequence analysis and neutralization sensitivity to paired maternal plasma

As maternal V3-specific IgG binding and tier 1 virus neutralizing responses were predictive of MTCT risk in this cohort [20], we explored possible signatures of neutralization resistance to paired maternal plasma in the V3 region. We examined the highly variable N and C-terminal region amino acid residues K305Q, I307T, H308T, R315Q, F317L, A319T, and D322R (Fig. 7), 3 of which (K305Q, I307T, and H308T) have previously been identified critical targets of the V3-specific IgG responses associated with reduced MTCT risk [43]. Comparing non-transmitted maternal sequences and infant T/F viruses, we found position K305R to be significantly associated with higher sensitivity to paired maternal plasma (p<0.001 by 1-sided permutation test). At position 308, sequences carrying mutations from the consensus amino acid H (N, P, S or T) were significantly associated with higher neutralization resistance to paired maternal plasma (p<0.001 by 1-sided permutation test). However, as for the MPER residues, we observed that these amino acids were equally distributed across non-transmitted maternal and infant T/F virus sequences, indicating that none of these amino acid residues were directly involved with transmission risk. This discrepancy could partially be due to the heterogeneity at amino acid residue position 308. Yet, 6 out of 14 transmitting mother-infant pairs exhibited distinct amino acid residues at position 308 from the more frequently occurring histidine (Fig. 7), suggesting variability at this position could be overrepresented in transmitting pairs.

## Discussion

While maternal and infant ART has considerably reduced rates of MTCT, pediatric HIV infection remains a significant public health problem in areas of high HIV prevalence, with up to 16% of HIV-infected women still transmitting the virus to their infant globally [1]. It is likely that a maternal or infant HIV-1 vaccine will be required to eliminate pediatric HIV [44]. However, a better understanding of factors that may drive the genetic bottleneck of virus populations in the setting of MTCT of HIV will be required to develop vaccination strategies that can block HIV transmission. Recent findings published by our group demonstrated that maternal V3 loop-specific and tier 1 virus-neutralizing antibody responses both correlated and were independently associated with of reduced MTCT risk. Moreover, we established that V3-specific antibodies in maternal plasma could neutralize maternal autologous viral variants circulating in plasma [20, 27]. To examine the potential role of maternal Env-specific responses in driving the viral genetic bottleneck of MTCT, we aimed to define if neutralization resistance to maternal autologous virus neutralizing antibodies is a defining feature of infant T/F variants compared to circulating maternal non-transmitted plasma variants.

In this cohort, 6 out of 16 (37%) peripartum-infected infants were infected by at least 2 T/F viruses. Interestingly, infection with multiple T/F viruses occurs in approximately 19-24% of heterosexual HIV infections [24, 45, 46], and 12-38% of homosexual infections [47-49], whereas up to 60% of infections that occur through intravenous drug use involve multiple T/Fs [50]. Thus, the rate of multiple T/F transmissions in this mother-infant cohort is in line with or slightly higher than sexual transmission modes, but lower than that of transmission via intravenous drug use. While it is well established that a genetic bottleneck occurs in the setting of MTCT, the determinants that drive the selection of 1 or multiple T/F viruses are less clear [4]. Importantly, the lack of maternal ART prophylaxis around the time of delivery in the WITS cohort could contribute to the observed high rate of multiple T/F viruses, potentially stemming from a larger virus inoculum in this cohort compared to ARV-treated mothers. Regardless, of the impact of maternal ART on the bottleneck of infant T/F viruses, maternal or infant immunization strategies will likely need to generate Env-specific responses that can block the diverse pool of maternal viruses circulating in plasma. Notably, these infant T/F viruses uniformly represented a minor variant of the maternal viral population Env variants, indicating that maternal antibodies that can block infant virus transmission will need to target minor circulating variants.

A greater understanding of virologic characteristics of infant T/F viruses will also be important to developing immune-based strategies to prevent MTCT. As maternal V3-specific IgG responses predicted reduced risk of transmission in this cohort, we investigated V3 loop residues in the maternal and infant viruses and how they related to paired maternal plasma neutralization sensitivity. Despite the association between maternal V3-specific IgG responses targeting the C terminal region and reduced MTCT risk in this cohort [43], we did not find amino acid residues within the C terminal region to be associated neutralization resistance to paired maternal plasma. Instead, we found that maternal non-transmitted and infant T/F viruses carrying N, P, S or T at the N terminal region amino acid residue position 308 were more neutralization resistant to paired maternal plasma. These seemingly disparate findings could partly be explained by several reasons. Firstly, N terminal amino acid residues 308 and 309 have been shown to interact with C terminal amino acid residue 317, and this interaction leads to the stabilization of the V3 loop [51, 52]. Thus, the disruption of intra-peptide interactions at either the N and or C terminal region could lead to altered neutralization sensitivity of viruses to paired maternal V3-specific IgG plasma responses. Secondly, it should be noted that in this study, we compared maternal non-transmitted circulating viruses to infant T/F viruses in 16 transmitting mother infant pairs, whereas we previously defined the potentially-protective role of maternal V3-specific IgG binding and neutralizing responses by comparing transmitting and non-transmitting women in the larger (n=248) WITS cohort [20, 43]. Finally, V3 loop accessibility to maternal neutralizing antibodies may be modulated by amino acid residues by distal amino acid residues within gp120 or gp41 [15]. For example, specific glycosylation sites within the V1 loop may alter V3 loop accessibility to V3-specific neutralizing antibodies [53]. Moreover, interactions between C2 and V3 may stabilize the structure of the HIV-1 Env [54], as demonstrated with the recent elucidation of the SOSIP trimer [55].

In contrast to previous studies that examined the neutralization sensitivity of randomly selected or non-paired infant or maternal virus isolates, our study defined the neutralization sensitivity of paired infant T/F viruses and maternal non-transmitted variants. Moreover, our study accounted for phylogenetic relationships of infant T/F viruses and maternal non-transmitted variants to represent the diverse maternal virus lineage pools. Furthermore, our study carefully controlled for analysis confounders such as transmission mode, disparate maternal and infant sample testing. Moreover, as the WITS cohort was enrolled and followed prior to the availability of ART to prevent MTCT, virus variant selection in this cohort is not influenced by ART selection pressures. With this robust study design, our analysis demonstrated that infant T/F viruses are mostly resistant to concurrent autologous maternal plasma, suggesting that infant T/F viruses are defined by neutralization resistance to maternal autologous virus neutralizing antibodies. This work confirms previous studies that have made this prediction based on smaller studies or with less well-defined maternal and infant virus variants [16, 56, 57]. Yet, Miligan *et. al* [58] recently showed that neutralization resistant viruses do not predict MTCT risk in a breastfeeding transmission setting. However, as our analysis focused on peripartum transmission only, there may be distinct virologic or immunologic determinants in peripartum and postpartum HIV transmission. Yet an important, a novel observation gleaned this study is that infant T/F viruses’ neutralization resistance to maternal plasma is not predictive of neutralization resistance to heterologous plasma. Remarkably, the tiered categorization of infant T/F viruses ranged from easy to neutralize tier 1 a viruses, to very difficult to neutralize tier 3 viruses, suggesting that heterologous plasma neutralization resistance is not a defining feature of infant T/F viruses. Specifically, 24% of infant T/F viruses isolated in this study were classified as “easy to neutralize” tier 1b or tier 1a variants by a standard panel of heterologous plasma [40], consistent with the hypothesis that these infant T/F viruses may be specifically resistant to maternal antibodies that co-evolved with the transmitted variants.

Not unsurprisingly, the majority of infant T/F viruses were neutralization sensitive to a number of second generation broadly neutralizing antibodies (Fig. 4). This finding is clinically relevant, as it suggests that infant passive immunization with second generation broad and potent bNAbs to prevent HIV-1 transmission could be an effective strategy to block MTCT. Interestingly, there is an ongoing passive immunization clinical trial of high-risk, HIV-exposed infants with VRC01 (https://clinicaltrials.gov/ct2/show/record/NCT02256631). The uniform sensitivity of these clade B infant T/F viruses isolated in our study to VRC01 neutralization suggests that these viruses would be effectively neutralized by VRC01, suggesting that clade B infant virus transmission may be blocked by VRC01.

To our knowledge, this is the largest study that has characterized infant T/F and maternal viruses and their neutralization sensitivity to maternal autologous virus neutralizing responses. Our study specifically addresses whether infant T/F viruses are defined by their neutralization sensitivity to maternal autologous virus neutralizing antibodies in peripartum MTCT of HIV. MTCT is a unique setting in which protective antibodies only need block autologous virus variants circulating in blood to which the infant is exposed. The observation that infant T/F viruses are neutralization resistant compared to non-transmitted maternal variants suggests that the development of a maternal vaccine that boosts maternal autologous virus neutralizing responses may be a viable strategy to further reduce MTCT risk. Maternal Env immunization regimens with closely related, but not identical, Envs to maternal circulating virus populations may elicit antibodies that target her autologous virus pool through the well-described immune phenomenon of ‘original antigenic sin’ [59, 60]. Our central finding that maternal autologous virus neutralization shapes the genetic bottleneck of peripartum transmission has important implications in designing maternal Env vaccination strategies that can synergize with current maternal ART treatment strategies help achieve an HIV-free generation.

## Materials and Methods

### Study Subjects and sample collection

Maternal and infant pairs from the WITS cohort that met the following criteria were selected: peripartum transmission, infant plasma samples from < 2.5 months of age, and maternal samples available from around delivery. Peripartum transmission was defined by negative a negative PCR result or negative culture from peripheral blood samples collected within 7 days of birth with subsequent a positive result 7 days after birth (Table 1).

**Table 1.** Maternal and infant pair clinical characteristics around delivery.

| Maternal and Infant ID | Year | Maternal Visit of sample collection | Mode of Transmission | Maternal plasma Viral Load (copies/mL) | Maternal Peripheral CD4+ T-cell count (cells/mm<sup>3</sup>) | Mode of Delivery | Weeks Gestation | Infant Age at sample Collection (days) | Infant Viral Load (copies/mL) | Infant Peripheral CD4+ T-cell count (cells/mm<sup>3</sup>) |
| --- | --- | --- | --- | --- | --- | --- | --- | --- | --- | --- |
| 100002 | 1993 | Delivery | Peripartum | 305,213 | 221 | Vaginal | 38 | 16 | 1,230,000 | 3326 |
| 100014 | 1993 | Delivery | Peripartum | 21,918 | 692* | C-section | 31 | 66 | 120,000 | 2012 |
| 100046 | 1993 | Delivery | Peripartum | 12,635 | 707 | Vaginal | 40 | 63 | 1,025,458 | 2509 |
| 100052 | 1993 | Delivery | Peripartum | 23,750 | 450 | Vaginal | 40 | 34 | 333,142 | 3219 |
| 100155 | 1992 | Delivery | Unknown | 87,193 | 318 | C-Section | 36 | 60 | NA | 3609 |
| 100307 | 1991 | Delivery | Unknown | 139,053 | 467* | C-section | 36 | 32 | 905,079 | 2308 |
| 100383 | 1991 | Delivery | Peripartum | 358,602 | 244 | Vaginal | 37 | 74 | 338,000 | 2071<sup>‡</sup> |
| 100504 | 1993 | NA | Unknown | 68287 | 1049 | C-Section | 39 | 30 | NA | 2742 |
| 100711 | 1990 | Delivery | Unknown | 4,104* | 571* | Vaginal | 39 | 64 | 341,691 | 1872<sup>‡</sup> |
| 100890 | 1991 | Delivery | Peripartum | 368,471 | 409 | Vaginal | 39 | 26 | 192,000<sup>‡</sup> | 2598 |
| 100997 | 1991 | Delivery | Peripartum | 175,526 | 413 | Vaginal | 38 | 20 | 46,289<sup>‡</sup> | 7048 |
| 101421 | 1991 | Delivery | Peripartum | 17,370 | 760 | Vaginal | 37 | 29 | 48,783 | 7628 |
| 101984 | 1991 | Delivery | Peripartum | 94,087 | 107 | Vaginal | 36 | 30 | 317969 | 1980 |
| 102149 | 1993 | Delivery | Unknown | 253,906 | 373 | Vaginal | 34 | 33 | 2,042,124 | 3181 |
| 102407 | 1993 | Delivery | Peripartum | 104,922 | 422 | Vaginal | 40 | 35 | 11,100 | 4677 |
| 102605 | 1992 | NA | Unknown | 5684 | 337 | Vaginal | 39 | 60 | 156639 | 1360 |

<sup>159</sup> Peripheral CD4+ T cell count and/or viral load reported from next available visit, within 2 months
<sup>160</sup> from the sample that was obtained for sequencing. NA =Not Available.
<sup>161</sup> <sup>‡</sup>Peripheral CD4+ T cell count and/or viral load reported from 25 weeks gestation, as CD4+ T cell
<sup>162</sup> count and/or viral load was not available at time of delivery.

### Ethics Statement

Samples used in this study were obtained from an existing cohort named as Women Infant Transmission Study (WITS). WITS cohort samples were received as de-identified material and were deemed as research not involving human subjects by Duke University Institutional Review Board (IRB). The reference number for that protocol and determination is Pro00016627.

### Viral RNA Extraction and SGA isolation

Viral RNA was purified from the plasma sample from each patient by the Qiagen QiaAmp viral RNA mini kit and subjected to cDNA synthesis using 1X reaction buffer, 0.5 mM of each deoxynucleoside triphosphate (dNTP), 5 mM DTT, 2 U/mL RNaseOUT, 10 U/mL of SuperScript III reverse transcription mix (Invitrogen), and 0.25 mM antisense primer 1.R3.B3R (5’-ACTACTTGAAGCACTCAAGGCAAGCT TTATTG-3’), located in the *nef* open reading frame. The resulting cDNA was end-point diluted in 96 well plates (Applied Biosystems, Inc.) and PCR amplified using Platinum Taq DNA polymerase High Fidelity (Invitrogen) so that < 30% of reactions were positive in order to maximize the likelihood of amplification from a single genome. A second round of PCR amplification was conducted using 2μl of the first round products as template. 07For7 (5’-AAATTAYAAAAATTCAAAATTTTCGGGTTTATTACAG-3’) and 2.R3.B6R (5’- TGA AGCACTCAAGGCAAGCTTTATTGAGGC -3’) were used as primer pair in the first round of PCR amplification step, followed by a second round with primers VIF1 (5’- GGGTTTATTACAGGGACAGCAGAG -3’)(nt 5960–5983 in the HXB2 tat coding region) and Low2c (5’- TGAGGCT TAAGCAGTGGGTT CC -3’) (nt 9413–9436 in HXB2 nef). PCR was carried out using 1X buffer, 2 mM MgSO4, 0.2 mM of each dNTP, 0.2μM of each primer, and 0.025 U/μl Platinum Taq High Fidelity polymerase (Invitrogen) in a 20μl reaction. Round 1 amplification conditions were 1 cycle of 94°C for 2 minutes, 35 cycles of 94°C for 15 seconds, 58°C for 30 seconds, and 68°C for 4 minutes, followed by 1 cycle of 68°C for 10 minutes. Round 2 conditions were one cycle of 94°C for 2 minutes, 45 cycles of 94°C for 15 seconds, 58°C for 30 seconds, and 68°C for 4 minutes, followed by 1 cycle of 68°C for 10 minutes. Round 2 PCR amplicons were visualized by agarose gel electrophoresis and sequenced for envelope gene using an ABI3730xl genetic analyzer (Applied Biosystems). The final amplification 3’-half genome product was ∼4160 nucleotides in length exclusive of primer sequences and included all of *rev* and *env* gp160, and 336 nucleotides of *nef*. Partially overlapping sequences from each amplicon were assembled and edited using Sequencher (Gene Codes, Inc). Sequences with double peaks per base read were discarded. Sequences with one double peak were retained as this most likely represents a Taq polymerase error in an early round of PCR rather than multiple template amplification; such sequence ambiguities were read as the consensus nucleotide. Sequence alignments and phylogenetic trees were constructed using ClustalW and Highlighter plots were created using the tool at https://www.hiv.lanl.gov/content/sequence/HIGHLIGHT/highlighter_top.html.

### Sequence Alignment

All maternal and infant envelope sequences were aligned using the Gene Cutter tool available at the Los Alamos National Laboratory (LANL) website (http://www.hiv.lanl.gov/content/sequence/GENE_CUTTER/cutter.html) and then refined manually. Full-length envelope sequences were manually trimmed in Seaview [61]. The infant T/F env virus sequences were visually identified looking at phylogenetic trees and highlighter plots, and infant consensus sequences of the major T/F lineage were created using the LANL Consensus Maker tool (http://www.hiv.lanl.gov/content/sequence/CONSENSUS/consensus.html). For infants that were infected by 2 or more distinct T/F viruses, the highlighter plots and phylogenetic trees were rooted on the consensus of the major variant.

### Infant T/F Virus Envelope Characterization

Maternal and infant envelope alignments were characterized using Bio-NJ phylogeny (Mega 6 Software) and highlighter plot (http://www.hiv.lanl.gov/content/sequence/HIGHLIGHT/HIGHLIGHT_XYPLOT/highlighter.html). The number of infant T/F viruses was determined by visual inspection of both phylogenetic trees and highlighter plots of infant-maternal env sequence alignments. Hypermutation was also evaluated using the tool Hypermut (http://www.hiv.lanl.gov/content/sequence/HYPERMUT/hypermut.html). Sequences with significant hypermutation (p<0.1) were removed from the alignment and not included in further analysis. When a sample was found to be overall enriched for hypermutation [24], positions within the APOBEC signature context were removed (Table 2). All 16 infant infections were acute and we were able to time the infection using the Poisson Fitter method after removing putative recombinants and/or hypermutated sequences as described above. Days since infant infection were calculated using the Poisson Fitter tool (http://www.hiv.lanl.gov/content/sequence/POISSON_FITTER/pfitter.html) which estimates the time since infection based on the accumulation of random mutations from the most recent common ancestor (MRCA) [39]. For infants infected with 2 or more T/F viruses, only the major variant was analyzed to obtain the time since the infection. The defined mutation rate was 2.16e-5. Values were reported in days with a 95% confidence interval and a goodness-of-fit p-value.

**Table 2.**
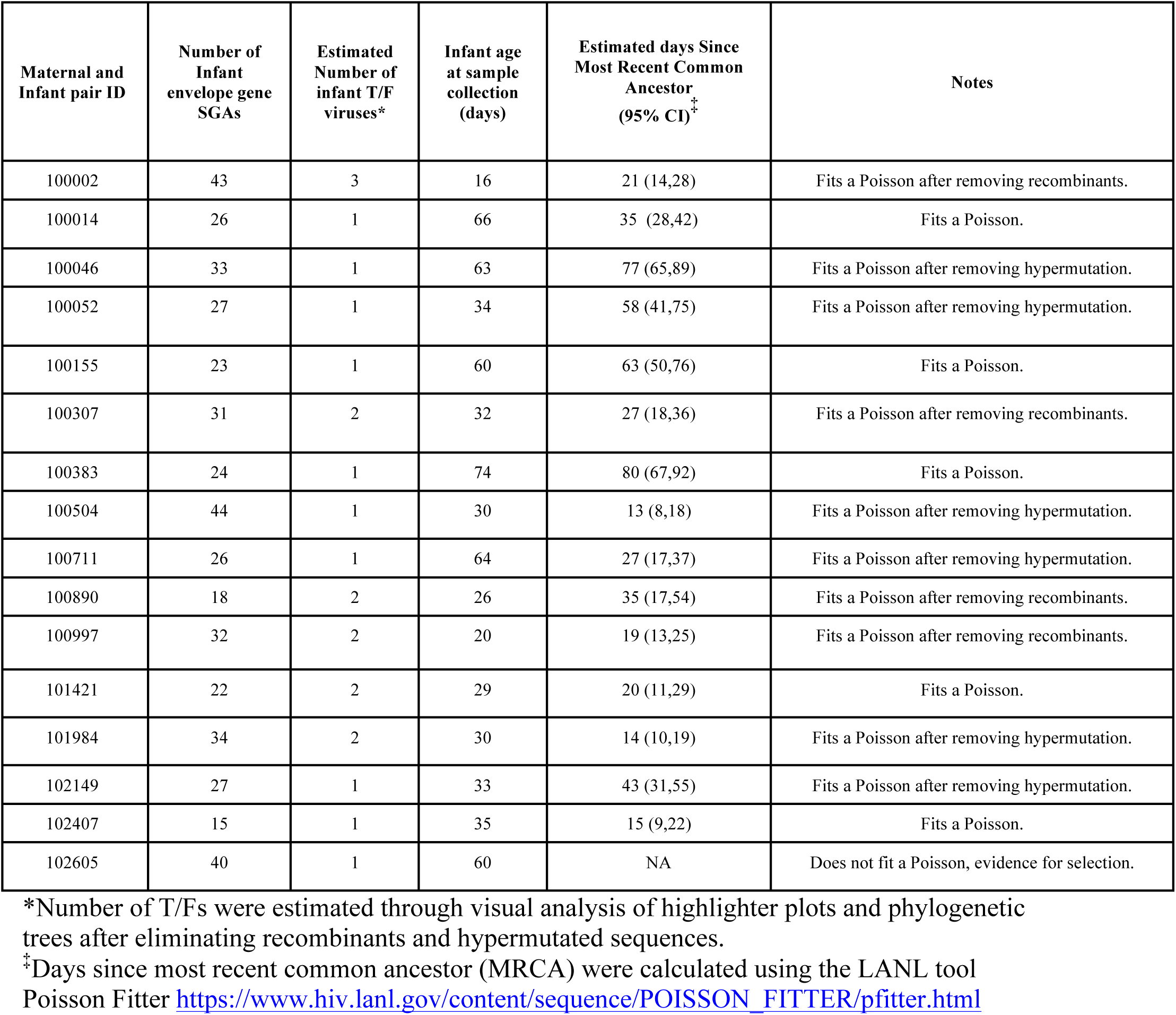
Number of sequences,T/F viruses,and estimated days since most common recent ancestor (MRCA) in infants.

### Infant T/F SGA Cloning

Amplicons from the first round PCR product that matched the infant consensus sequence (T/F virus sequence) were ligated into pcDNA3.1 Directional Topo vectors (Invitrogen) by introducing a –CACC 5’ end via a PCR reaction with the primers Rev19 (5’-ACTTTTTGACCACTTGCCACCCAT-3’) and Env1A (5’-caccTTAGGCATCTCCT ATGGCAGGAAGAAG-3’). Phusion^®^ High-Fidelity PCR Master Mix with HF Buffer was used according to the manufacturer’s instructions (New England BioLabs). Plasmids were then transformed into XL10 gold chemically competent Escherichia coli cells. Cultures were grown at 37°C for 16 hours. Colonies were selected for growth, and plasmids were minipreped and quality controlled by restriction enzyme digestion using BamHI and XhoI (New England BioLabs). Plasmids containing an insert of correct size were sequenced to confirm 100% sequence homology with the original env infant consensus sequence. Plasmids were then prepared by Megaprep (Qiagen) kit and re-sequenced to confirm. For three infants, 100046, 100383 and 101580, no single genome isolated matched 100% of nucleotides in the consensus sequence. Therefore site-directed mutagenesis on a single nucleotide was performed to create an isolate identical to the consensus sequence. Primers for site directed mutagenesis were designed using Agilent’s QuikChange primer design program and Agilent’s QuikChange II XL kit was used. Sequencing of the clones was done to ensure 100% homology with the infant consensus sequence.

### Pseudovirus preparation

Env pseudoviruses were prepared by transfection in HEK293T (ATCC, Manassas, VA) cells with 4μg of env plasmid DNA and 4μg of env-deficient HIV plasmid DNA using the FuGene 6 transfection reagent (Roche Diagnostics) in a T75 flask. Two days after transfection, the culture supernatant containing pseudoviruses was harvested, filtered, aliquoted, and stored at −80°C. An aliquot of frozen pseudovirus was used to measure the infectivity in TZM-bl cells. 20μl of pseudovirus was distributed in duplicate to 96-well flat bottom plates (Co-star). Then, freshly trypsinized TZM-bl cells were added (10,000 cells/well in Dulbecco’s modified Eagle’s medium (DMEM)-10% fetal bovine serum (FBS) containing HEPES and 10 *µ* g/ml of DEAE-dextran). After 48 h of incubation at 37°C, 100μl of medium was removed from the wells. 100μl of luciferase reagent was added to each well and incubated at room temperature for 2 min. 100μl of the lysate was transferred to a 96-well black solid plate (Costar), and the luminescence was measured using the Bright-Glo™ luminescence reporter gene assay system (Promega).

### Neutralization Assays

Neutralizing antibody activity was measured in 96-well culture plates by using Tat-regulated luciferase (Luc) reporter gene expression to quantify reductions in virus infection in TZM-bl cells. TZM-bl cells were obtained from the NIH AIDS Research and Reference Reagent Program, as contributed by John Kappes and Xiaoyun Wu. Assays were performed with HIV-1 Env-pseudotyped viruses as described previously [62]. Test samples were diluted over a range of 1:20 to 1:43740 in cell culture medium and pre-incubated with virus (∼150,000 relative light unit equivalents) for 1 hr at 37°C before addition of cells. Following a 48 hr incubation, cells were lysed and Luc activity determined using a microtiter plate luminometer and BriteLite Plus Reagent (Perkin Elmer). Neutralization titers are the sample dilution (for serum/plasma) or antibody concentration (for sCD4, purified IgG preparations and monoclonal antibodies) at which relative luminescence units (RLU) were reduced by 50% compared to RLU in virus control wells after subtraction of background RLU in cell control wells. Serum/plasma samples were heat-inactivated at 56°C for 1 hr prior to assay. Murine leukemia virus SVA.MLV was used as a negative control [40]. A response was considered positive if the plasma ID_50_ against infant T/F viruses was at least 3 times higher than the ID_50_ versus SVA.MLV.

### Env Virus Variant Tier Phenotyping Assay

Neutralization titers (ID50s) were determined essentially as described above using five plasma samples from HIV+ individuals in chronic infection. The geometric mean titer (GMT) was calculated in Microsoft Excel and tier phenotype was determined by comparing these values to the GMTs of standard panels of viruses representing tier 1A, tier 1B and tier 2 viruses [40, 63] using the same five HIV+ plasma samples.

### Sequence Selection Algorithm

To select maternal non-transmitted variants and capture the most divergent sequences from the infant T/F, we devised an algorithm as follows. The algorithm finds the most variable positions in the amino acid alignment and ranks all sequences with respect to the frequencies at these positions. Sequences are then selected starting from the most divergent based on motif coverage as observed in the alignment and in the phylogenetic tree (in other words, if a group of diverging sequences all share the same motif, only one in the group and/or tree node is selected).

### Statistical Analysis

To test whether infant transmitted viruses were statistically significantly more resistant to maternal plasma than non-transmitted maternal sequences, we devised a 1-sided permutation test. At each iteration, we randomly assigned the “transmitted” status to any one sequence in each infant-mother pair, and then ranked the remaining sequences in the pair according to maternal plasma responses. All ranks across all pairs were then summed. We repeated this randomization 1,000 times and then calculated the p-value as the percentage of sum of ranks that were above the observed sum of ranks, out of all randomizations performed. This method is robust, as it does not make any underlying assumption of the distribution of the maternal plasmas, and it preserves the within mother-infant correlation of the data. The same algorithm was used to test whether specific amino acid positions conferred resistance to maternal plasma and/or antibodies. This time the “transmitted” status that was reshuffled at each iteration was the wild type amino acid.

## Acknowledgements

We acknowledge Nathan Vandergrift in selecting peripartum transmission samples from the WITS cohort.

## Supporting Information

**Fig. S1 Phylogenetic tree and highlighter plot for all other 12 mother infant pairs.** Infant sequences are labeled in red circles and maternal sequences are labeled in blue squares.

**Fig. S2 Phylogenetic tree and highlighter plot for mother infant pair 102605.** Infant sequences are labeled in red and maternal sequences are labeled in blue. The non-random accumulation of synonymous mutations in the infant (which caused the Poisson Fitter analysis to fail) is evident on the right as marked by red box.

**Fig. S3 Neutralization sensitivity of Infant T/F viruses to heterologous broadly neutralizing antibodies (bNAbs).** Dark colors represent easily neutralized viruses. Second T/F viruses are marked with an *.

